# Dysregulated TDP-43 proteostasis perturbs excitability of spinal motor neurons during brainstem-mediated fictive locomotion in zebrafish

**DOI:** 10.1101/2023.05.12.540624

**Authors:** Kazuhide Asakawa, Hiroshi Handa, Koichi Kawakami

## Abstract

Spinal motor neurons (SMNs) are the primary target of degeneration in amyotrophic lateral sclerosis (ALS). Degenerating motor neurons accumulate cytoplasmic TAR DNA-binding protein 43 (TDP-43) aggregates in most ALS cases. This SMN pathology can occur without mutation in the coding sequence of the TDP-43-encoding gene, TARDBP. Whether and how wild-type TDP-43 drives pathological changes in SMNs in vivo remain largely unexplored. In this study, we develop a two-photon calcium imaging setup in which tactile-evoked neural responses of motor neurons in the brainstem and spinal cord can be monitored using the calcium indicator GCaMP. We devise a piezo-assisted tactile stimulator that reproducibly evokes a brainstem descending neuron upon tactile stimulation of the head. A direct comparison between caudal primary motor neurons (CaPs) with or without TDP-43 overexpression in contiguous spinal segments demonstrates that CaPs overexpressing TDP-43 display attenuated Ca^2+^ transients during fictive escape locomotion evoked by the tactile stimulation. These results show that excessive amounts of TDP-43 protein reduce the neuronal excitability of SMNs and potentially contribute to asymptomatic pathological lesions of SMNs and movement disorders in patients with ALS.

## INTRDUCTION

Amyotrophic lateral sclerosis (ALS) is a neurodegenerative disorder characterized by the progressive loss of both upper and lower motor neurons. A hallmark of ALS is the deposition of cytoplasmic aggregates of TAR DNA-binding protein of 43 kDa (TDP-43) in degenerating motor neurons (Arai et al., 2006; Neumann et al., 2006). Cytoplasmic TDP-43 aggregates have been detected in almost all cases of sporadic ALS (Mackenzie et al., 2007), accounting for more than 90% of all ALS cases (Taylor, Brown, & Cleveland, 2016). The remaining 10% of ALS cases are heritable forms of ALS, in which mutations in the TDP-43/TARDBP gene are identifiable at a certain frequency.

An important but largely unanswered question in ALS is whether and how dysregulation of wild-type TDP-43 affects function of motor neurons in sporadic ALS cases, in which mutations in the coding sequence of the TDP-43/TARBDP gene are absent. TDP-43 is an RNA/DNA-binding protein with wide range of target nucleic acids, and its dysfunction likely has a widespread impact on cellular transcriptome and proteome (Acharya, Govind, Shore, Stoler, & Reddi, 2006; Li et al., 2022; Ou, Wu, Harrich, Garcia-Martinez, & Gaynor, 1995; Polymenidou et al., 2011; Sephton et al., 2011; Tollervey et al., 2011; Xiao et al., 2011). Recent studies have revealed that a significant consequence of the loss of TDP-43 function is the occurrence of splicing abnormalities, leading to erroneous incorporation of intronic sequences into mature mRNA (Brown et al., 2022; Koike et al., 2023; Ling, Pletnikova, Troncoso, & Wong, 2015; Ma et al., 2022; Mehta, Brown, Ward, & Fratta, 2023). Overexpression of wild-type TDP43 can also be toxic in various models (Asakawa, Handa, & Kawakami, 2020; Barmada et al., 2010; Estes et al., 2011; Johnson, McCaffery, Lindquist, & Gitler, 2008; Kabashi et al., 2010; Wils et al., 2010). Excessive amounts of wild-type TDP-43 impair splicing accuracy. For example, overexpression of wild-type TDP-43 in human cultured cells causes exon skipping in the pre-mRNA of the cystic fibrosis transmembrane conductance regulator (CFTR) protein, resulting in an inactive translation product (Buratti et al., 2001). The effect of the molecular-level dysfunction of wild-type TDP-43 on motor neuron excitability is poorly understood. In the mouse model, ectopic expression of the human nuclear localization sequence-deficient TDP-43 (TDP-43ΔNLS) results in hyperexcitability of layer V excitatory neurons in the cortex, while that of the human wild type TDP-43 is tolerated only with subtle electrophysiological changes in the cortex (Dyer et al., 2021). In the zebrafish model, injection of mRNA encoding a human mutant TDP-43 at the one-cell stage blastula results in the augmentation of spinal cord glutamatergic synaptic currents in primary motor neurons in the spinal cord; however, no discernible phenotype has been described from the overexpression of the wild-type TDP-43 protein (Petel Legare et al., 2019).

In this study, we investigated the effects of excessive TDP-43 protein levels on the neural activity of spinal motor neurons (SMNs) in the context of escape locomotion in zebrafish larvae. Larval zebrafish provide a valuable model for studying motor neuron function because of their translucent bodies, allowing for direct visualization of motor neurons in the spinal cord at single-cell resolution. We utilized the genetically encoded calcium imaging probe GCaMP7a to monitor the neural activity of SMNs during tactile-induced fictive escape locomotion and found that SMNs overexpressing fluorescently tagged wild-type TDP-43 displayed reduced calcium transients during fictive escape locomotion. These results suggest that excessive amounts of TDP-43 protein reduce the neuronal excitability of SMNs during locomotion, potentially contributing to movement disorders in ALS.

## MATERIALS AND METHODS

### Fish husbandry

This study was performed in accordance with the Guide for the Care and Use of Laboratory Animals of the Institutional Animal Care and Use Committee of the National Institute of Genetics (NIG, Japan). Fish were raised under 12:12 light/dark (L/D) cycles during the first five days after birth.

### Tg[vglut2a:Gal4] line

For generation of the BAC transgene Tg[vglut2a:Gal4], the BAC clone DKEY-145P24 was used. The *hsp70l* promoter (650 bp)-Ga4FF-polyA-Km^r^ cassette was introduced downstream of the vglut2a 5’-UTR in the vglut2a-BAC DNA in SW102 cells (Asakawa & Kawakami, 2018; Warming, Costantino, Court, Jenkins, & Copeland, 2005). For transgenesis of the engineered vglut2a-BAC, the iTol2-amp cassette (Suster, Abe, Schouw, & Kawakami, 2011) was amplified by PCR with the primer pair iTol2A-IndB5-f (5’-cga gcc gga agc ata aag tgt aaa gcc tgg gg tgc cta atg agt gag cta CCC TGC TCG AGC CGG GCC C-3’) and iTol2A-IndB5-r (5’-ggt ttc ccg act gga aag cgg gca gtg agc gca acg caa tta atg tga gtA TTA TGA TCC TCT AGA TCA GAT CT-3’), where the lower and upper cases are pIndigoBAC-5 sequences for homologous recombination and iTol2-amp annealing sequences, respectively, and was introduced into the backbone (pIndigoBAC-5, GenBank Accession; EU140754). The resulting vglut2a-BAC carrying the iTol2-amp cassette was purified and injected into one-cell stage zebrafish embryos with Tol2 transposase mRNA (Asakawa, Abe, & Kawakami, 2013).

### Kaede photoconversion

The trigeminal ganglion of Tg[vglut2a-Gal4] Tg[vglut2a:Gal4] Tg[UAS:Kaede] fish embedded in 1% low-melt agarose was irradiated with a laser at a wavelength of 405 nm using a confocal laser scanning microscope (FV-1200D, Olympus) at 44 hours post-fertilization (hpf).

### Calcium imaging

Two-photon calcium imaging was performed using LSM7MP (Zeiss) equipped with MaiTai-eHPDS (SpectraPhysics). To minimize body movement during imaging, fish at 48 hpf were treated with a neuromuscular junction blocker, d-tubocurarine (100 μM, Sigma; T2379), after their tail tips were cut and embedded in 2% low-melting agarose (Ronza). The agarose around the head region was removed for tactile stimulation using a tungsten needle (NILACO) attached to an electrode holder (H-12, NARISHIGE) controlled by a piezo actuator (MC-140L, MESS-TEK), piezo-driver (M-2674, MESS-TEK), and function generator (DF-1906).

### Data analysis

Data analysis was performed using the Fiji/ImageJ software. Statistical analysis was performed with using GraphPad Prism Software.

## RESULTS AND DISCUSSION

To investigate the effect of dysregulated TDP-43 proteostasis on SMNs in vivo, we focused on the brainstem-mediated escape locomotor circuit that assembles during the larval stage of zebrafish. Zebrafish develops neural circuits for robust escape locomotion against tactile stimuli by the time of hatching (approximately 48 hpf) (Saint-Amant & Drapeau, 1998). Tactile sensation in the head region is perceived through the peripheral sensory arbors of the trigeminal sensory neurons that tile the body surface (Sagasti, Guido, Raible, & Schier, 2005). The central axons of the trigeminal sensory neurons extend along the descending longitudinal fiber pathway and contact the brainstem descending neurons, including the Mauthner cell, which is activated during the escape response (Kimmel, Hatta, & Metcalfe, 1990). We explored the anatomical relationship between the trigeminal sensory neurons and the Mauthner cell at 48 hpf by simultaneously visualizing the central axon of the trigeminal sensory neurons and Mauthner cell by combining a bacterial artificial chromosome (BAC) transgenic Gal4 line for vglut2a (L-glutamate transmembrane transporter) (Fig.1a), the Gal4 driver for the Mauthner cell Tg[hspGFFDMC130A] (Asakawa et al., 2008; Pujol-Marti et al., 2012), and Tg[UAS:Kaede] (Scott et al., 2007). The central axons of the trigeminal sensory neurons were highlighted with red fluorescence by the photoconversion of Kaede, which was expressed the trigeminal ganglion, with a laser light with a 405 nm wavelength in the Tg[vglut2a-Gal4] Tg[vglut2a:Gal4] Tg[UAS:Kaede] triple transgenic fish at 48 hpf (Fig. 1b). The central axons of the trigeminal sensory neurons descended in the vicinity of the lateral region of the Mauthner cell soma (Fig. 1b), suggesting neuronal contact between the bundle of central axons of the trigeminal sensory neurons and the Mauthner cell, consistent with the previous observations (Kimmel et al., 1990). To establish a functional connection between trigeminal sensory neurons and Mauthner cells at 48 hpf, we set up a system that enables the tactile-evoked neuronal activity of Mauthner cell to be visualized by calcium imaging under two-photon microscopy. The Tg[hspGFFDMC130A] driver was crossed with Tg[UAS:GCaMP7a] (Muto, Ohkura, Abe, Nakai, & Kawakami, 2013) to express the calcium indicator GCaMP7a in Mauthner cells. Tg[hspGFFDMC130A] Tg[vglut2a:Gal4] Tg[UAS:GCaMP7a] triple transgenic fish at 48 hpf were briefly treated with a neuromuscular blocker, embedded upright in agarose, and subjected to calcium imaging under a two-photon microscope. We set up a piezo-controlled tungsten needle for controlled application of tactile stimuli. A single z-section including the bilateral Mauthner cells was scanned using a two-photon laser at 2.5 Hz (Fig. 1c). When a tactile stimulus was applied to the head skin near the eye during imaging, the Mauthner cell ipsilateral to the stimulus strongly increased GCaMP7a fluorescence, whereas the contralateral Mauthner cell displayed only a minimal increase in the GCaMP7a signal (Fig. 1c, d). These observations demonstrate that trigeminal sensory neurons and ipsilateral Mauthner cell establish functional neural connections by 48 hpf and likely contribute to touch-evoked escape locomotion at this stage.

**Figure1.**
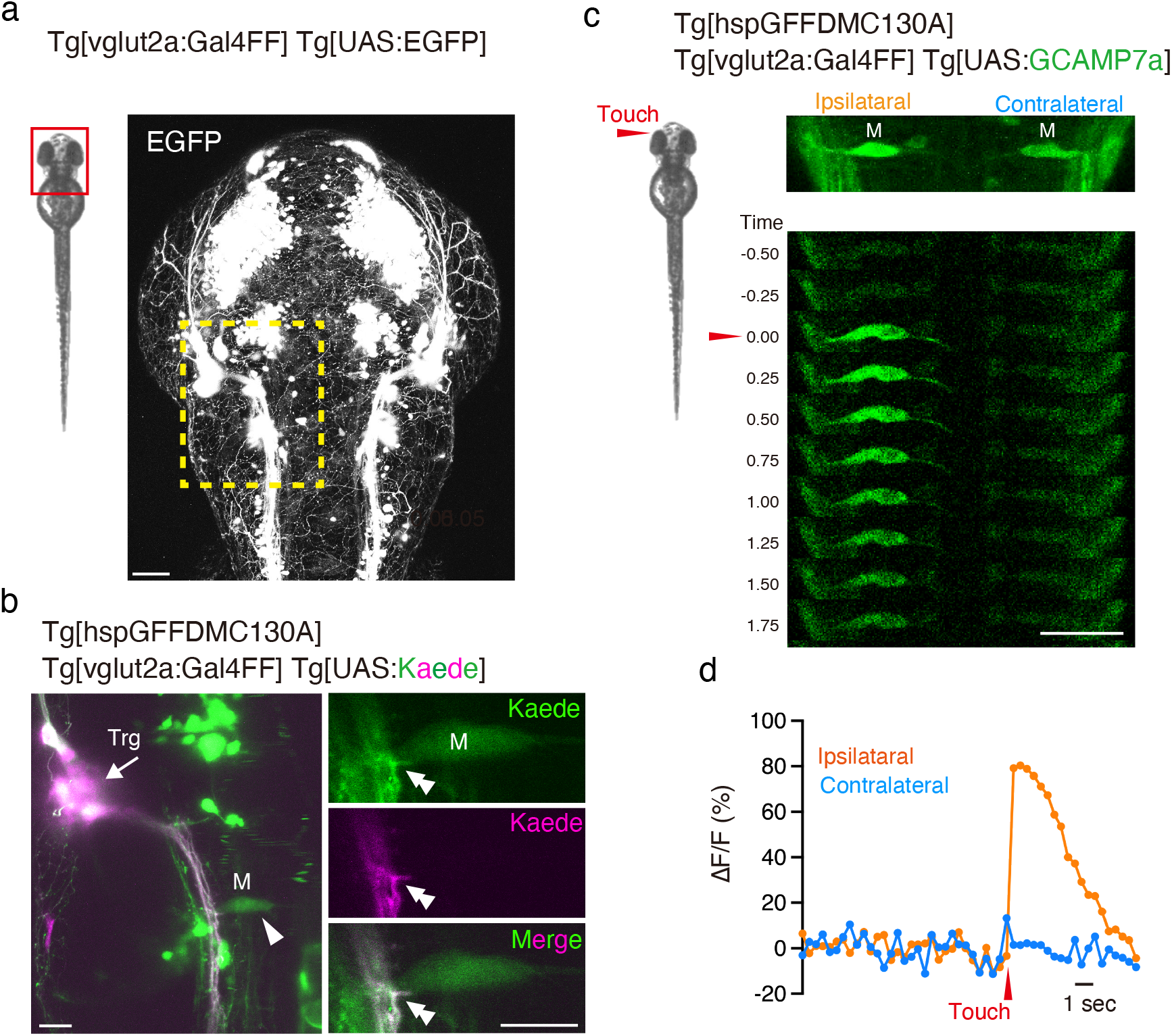
Anatomical and functional connections between trigeminal sensory neurons and Mauthner cells in zebrafish larvae a, Dorsal view of the head region of Tg[vglut2a:Gal4FF] Tg[UAS:EGFP] fish at 48 hpf. The dashed rectangle indicates the area corresponding to the brainstem region analyzed in b. b, Left panel: Kaede expressed in the trigeminal sensory neurons is photoconverted by illuminating the trigeminal ganglion region (Trg, arrow) with a laser light with a 405 nm wavelength (magenta). Afferent axons are projected towards a Mauthner cell (arrowhead) highlighted in magenta. Right panels: A single confocal section of the contact site (double arrowheads) of the afferent axons of trigeminal sensory neurons and Mauthner cells. c, Calcium imaging of the Mauthner cells before and after tactile stimulation. d, ΔF/F of the neural response of Mauthner cells in c. Scale bars indicate 50 μm (c) or 20 μm (a, b).

Having established a tactile stimulation setup under a two-photon microscope, we investigated the effect of wild-type TDP-43 overexpression on excitability of SMNs during tactile-induced fictive escape locomotion. We crossed the Tg[SAIG213A] Gal4 driver with Tg[UAS:GCaMP7a] to drive GCaMP7a expression in caudal primary motor neurons (CaPs), which innervates the ventral third of the myotome (Myers, Eisen, & Westerfield, 1986). To overexpress TDP-43 in CaPs, we used Tg[UAS:mRFP1-TDP-43z] line, which expresses mRFP1-tagged zebrafish TDP-43/tardbp in a Gal4-dependent manner that primarily localizes to the nuclei of CaPs (Asakawa et al., 2020). While, in Tg[SAIG213A] Tg[UAS:mRFP1-TDP-43z] Tg[UAS:GCaMP7a] triple transgenic fish, GCaMP7a was expressed in almost all CaPs, mRFP1-TDP-43z was not detectable in some of these CaPs, owing to transgene silencing (Fig. 2a). We took advantage of the variegated expression of mRFP1-TDP-43z and used CaPs expressing GCaMP7a without a nuclear mRFP1-TDP-43z signal as an internal control. We set a region of interest in the spinal cord of laterally positioned Tg[SAIG213A] Tg[UAS:mRFP1-TDP-43z] Tg[UAS:GCaMP7a] fish embedded in agarose and a single z-section that sliced two to three CaPs expressing GCaMP7a; at least one of which did not contain the nuclear mRFP1-TDP-43z signal. When a tactile stimulus was applied to the head, the GCaMP7a signals of the CaPs robustly increased and persisted for approximately 10 s, regardless of the presence of the nuclear mRFP1-TDP-43z signal (Fig. 2b). While a collective comparison of the maximum ΔF/F signals of GCaMP7a from all trials did not reveal a significant effect of mRFP1-TDP-43z expression (Fig. 2c), pairwise comparisons within adjacent spinal segments confirmed that mRFP1-TDP-43z overexpression significantly attenuated the maximum ΔF/F signal of the CaPs (Fig. 2d). Notably, the CaPs on both sides of the spinal hemisegments were activatable in response to lateralized tactile stimulation, suggesting that the ΔF/F signal of GCaMP7a in the CaPs reflects not only neural excitation via contralateral monosynaptic connection by the Mauthner cell (Jontes, Buchanan, & Smith, 2000), but also burst activity during fictive escape swimming. Taken together, these observations indicated that TDP-43 overexpression reduced the neural excitability of SMNs during locomotion.

**Figure2.**
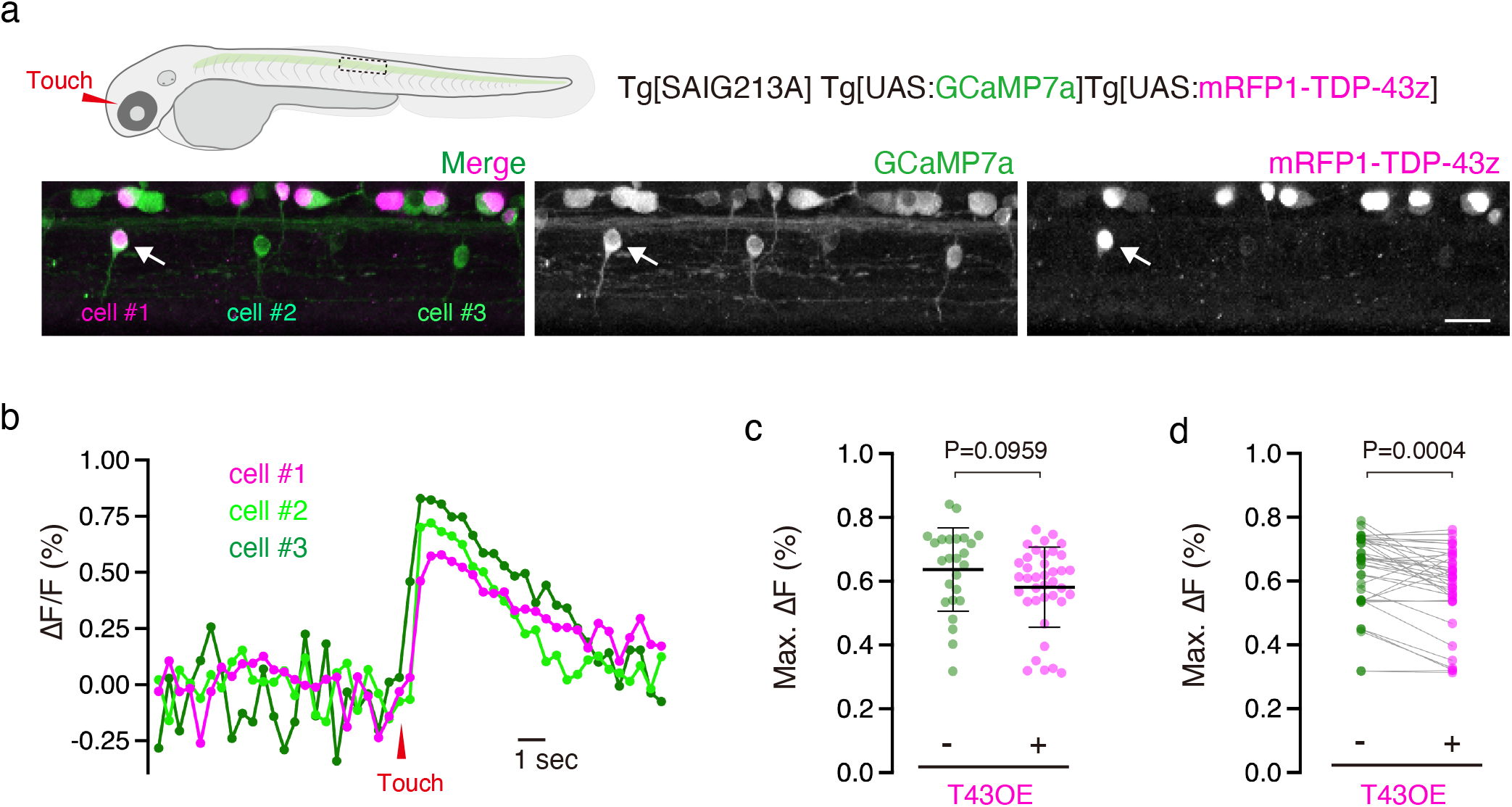
Calcium imaging of the CaPs overexpressing mRFP1-TDP-43z a Lateral view of the spinal cord of Tg[SAIG213A] Tg[UAS:GCaMP7a] Tg[UAS:mRFP1-TDP-43z] fish at 48 hpf. Among the three CaPs shown, cell #1 (arrow) displayed the nuclear mRFP1-TDP-43z signal, whereas cells #2 and cell #3 did not. b, ΔF/F of the three CaPs before and after tactile stimulation of the head. c,d, Comparison of maximum ΔF/F values between CaPs with or without nuclear mRFP1-TDP-43z signal in non-paired (c) and paired *t*-tests (d). Results were obtained from 38 trials of 14 CaPs with nuclear mRFP1-TDP-43z and 10 CaPs without nuclear mRFP1-TDP-43z in three animals. The scale bar indicates 20 μm.

In the present study, we performed two-photon calcium imaging of the neural activity of the escape locomotor circuit and found that the overexpression of mRFP1-tagged wild-type TDP-43 reduced the neural excitability of the SMNs during tactile-induced fictive escape locomotion. Reduced neural excitability has been associated with the overexpression of TDP-43 carrying ALS mutations in several animal models, but not with the overexpression of wild-type TDP-43 (Arnold et al., 2013; Dyer et al., 2021; Petel Legare et al., 2019). Direct comparison of the neural activity of SMNs with or without TDP-43 overexpression at a single-cell resolution clearly showed that wild-type TDP-43 affects the excitability of SMNs when in excess.

By definition, a decrease in ΔF/F indicates a reduction in the peak intracellular Ca^2+^ level by neural excitation when an excessive amount of TDP-43 is present in the SMNs. Several possible mechanisms may explain this ΔF/F decrease. The first is that the influx of extracellular Ca^2+^ into the cytosol upon plasma membrane depolarization is attenuated in CaPs overexpressing TDP-43. TDP-43 is known to regulate the transcription of voltage-gated Ca^2+^ channels (CaV1.2) in the pancreatic β-cell line MIN6, and TDP-43 depletion downregulates CaV1.2 at the transcriptional level (Araki et al., 2019). The physical association of TDP-43 with CaV1.2 mRNA was shown in the mouse brain by cross-linking and immunoprecipitation experiments (Polymenidou et al., 2011). Furthermore, locomotor deficits in flies lacking neuronal TDP-43 are rescued by exogenous expression of voltage-gated calcium channels (Chang, Hazelett, Stewart, & Morton, 2014). Locomotor defects in zebrafish expressing TDP-43 with the ALS mutation G348C were restored by treatment with L-type calcium channel agonists (Armstrong & Drapeau, 2013). These observations support the hypothesis that reduced Ca^2+^ influx via CaV1.2 causes attenuated neuronal excitability in SMNs overexpressing TDP-43. Another possibility that is not mutually exclusive with the first is that the insufficient release of Ca^2+^ from the endoplasmic reticulum (ER) reduces the level of cytosolic Ca^2+^ upon neural excitation. Overexpression, but not reduction of function, of TDP-43 reduces the binding of VAPB to PTPIP51, which is essential for the mitochondria-associated membrane (MAM) function, in both transfected cells and transgenic mice (Stoica et al., 2014). Given the critical role of MAM in intracellular Ca^2+^ homeostasis, TDP-43 overexpression-dependent MAM disruption may cause a reduction in ΔF/F. It is possible that other cellular modules involved in Ca^2+^ homeostasis and/or motor neuron excitability are affected by mRFP1-TDP-43z overexpression.

A limitation of this study is that we could not quantitatively determined the extent to which mRFP1-TDP-43z is overexpressed in CaPs in relation to endogenous TDP-43 proteins expressed from two paralogue genes for TDP-43 in zebrafish, *tardbp* and *tardbpl*. However, mRFP1-TDP-43z overexpression by the Tg[SAIG213A] driver did not lead to the deposition of mRFP1-TDP-43z aggregates in the cytoplasm of CaPs (Asakawa et al., 2020), showing that Tg[SAIG213A]-driven mRFP1-TDP-43z overexpression is excessive enough to cause cytotoxicity, but not sufficient to induce the formation of its cytoplasmic aggregates. We have previously shown that the Tg[SAIG213A]-driven mRFP1-TDP-43z overexpression halts the axon outgrowth of CaPs (Asakawa et al., 2020). The present study further demonstrates that the axon outgrowth defects is accompanied by reduced neural excitability. Thus, we posit that the reduced motor unit size and diminished neural excitability caused by mRFP1-TDP-43z overexpression might correspond to asymptomatic pathological lesions in SMNs that occur before the accumulation of cytoplasmic TDP-43 aggregates and later contribute to the manifestation of motor dysfunction in ALS.

## AUTHOR CONTRIBUTIONS

K.A. conceived the research and designed and performed the experiments. K.A. and K.K. analyzed the data. K.A., HH., and K.K. wrote the manuscript.

## ACKNOWLEDGMENTS

The authors thank the members of the Kawakami lab for their generous support. We are grateful to Akira Muto for the guidance on calcium imaging using GCaMP7a. This work was supported by the Nakabayashi Trust For ALS Research (K.A.), The Kato Memorial Trust For Nambyo Research (K.A.), Daiichi-Sankyo Foundation of Life Science (K.A.), Takeda Science Foundation (K.A.), KAKENHI grant numbers JP16K07045 (K.A.), JP19K06933 (K.A.), JP22H02958 (K.A.), JP23H04266 (K.A.), JP21H02463 (K.K.) and AMED-PRIME grant number JP23gm6410011h0003 (K.A.), and the National BioResource Project (NBRP) (K.K.).

## Notes

### Competing Interest Statement

The authors have declared no competing interest.

